# Differential Cerebral White Matter Tract Alterations in Generalized Anxiety Disorder Revealed by Ultra-High Field 7T Diffusion-Weighted Imaging

**DOI:** 10.64898/2025.12.26.696598

**Authors:** Chang-Le Chen, Minjie Wu, Yueyang Chi, Max Rose, Jessica C. Weber, Shannon T. Lamb, Hecheng Jin, Tamer S. Ibrahim, Cecile D. Ladouceur, Yue-Fang Chang, Kuei Y Tseng, Howard J. Aizenstein, Shaolin Yang

## Abstract

**Background:** Generalized anxiety disorder (GAD) is characterized by chronic worry and emotional dysregulation, yet its underlying white matter (WM) architecture remains inconsistent in previous neuroimaging studies. This study aimed to delineate microstructural WM alterations in GAD using ultra-high field (7T) diffusion tensor imaging (DTI) and advanced correlational tractography, evaluating their associations with symptom severity and diagnostic-aided value.

**Methods:** Eighty-eight participants (27 GAD, 61 healthy controls, HCs) underwent 7T DTI scanning with 1.5-mm isotropic resolution. Whole-brain correlational tractography was applied to identify tracts exhibiting significant group-related differences in diffusion indices while controlling for demographic covariates. Associations with Hamilton Anxiety Rating Scale (HAM-A) and Penn State Worry Questionnaire (PSWQ) scores were examined, and machine learning-based models were used to assess the diagnostic-aided utility of identified WM features.

**Results:** Two tract bundles showed significant fractional anisotropy (FA) alterations in GAD: decreased FA in the right prefrontal pathway (*P*_FDR_ = 0.039) and increased FA in the right cingulum (*P*_FDR_ < 0.001). The anterior portions of both tracts exhibited stronger effects of GAD. FA in the right cingulum positively correlated with HAM-A and PSWQ scores, indicating that greater microstructural coherence was associated with higher anxiety and worry severity (both *P*_FDR_’s < 0.001). Inclusion of WM features could improve classification performance beyond neuropsychological measures alone.

**Conclusions:** Ultra-high field 7T tractometry revealed a differential pattern of WM abnormalities in GAD, suggesting weakened prefrontal control and hyper-integrated cingulate connectivity as structural correlates of emotional dysregulation. These findings indicate the potential of 7T DTI markers for advancing mechanistic and diagnostic understanding of GAD.

## 1 Introduction

Generalized anxiety disorder (GAD) is a highly prevalent (6.2% lifetime prevalence) and debilitating mental illness (1). It is defined by chronic, excessive, and often uncontrollable worry that permeates multiple aspects of a person’s life, causing significant psychological distress and functional impairment (2). Critically, GAD is often poorly recognized in general clinical settings, leading to high rates of comorbidity (1). This diagnostic overlap obscures the disorder’s specific pathology and hampers the generalizability of neuroscientific research findings. Consequently, a deeper understanding of the specific neurobiological architecture underlying GAD is essential to enhance diagnostic reliability and facilitate the development of more precise and targeted therapeutic interventions.

One of the current cognitive frameworks positions GAD as a disorder of emotion dysregulation (3, 4). Neuroimaging studies have implicated dysfunctional communication within the cortico-limbic circuit, the primary neural system responsible for coordinating emotional processing and executive control (5, 6). Functional magnetic resonance imaging (fMRI) research has demonstrated difficulties in engaging the prefrontal cortex (PFC) and anterior cingulate cortex (ACC) during emotional regulation tasks, often interpreted as alterations in the top-down control system that normally inhibits limbic responses (4, 6, 7). However, the precise mechanisms underlying this cortical dysfunction remain the subject of competing hypotheses. One prominent view suggests that GAD involves hypoactive PFC/ACC function, leading to insufficient emotional regulation and unchecked amygdala reactivity (7, 8). Conversely, some fMRI studies reveal hyperactivation in prefrontal cortical areas during rest and task engagement, suggesting an overactive top-down control system associated with chronic worry (3, 9). These aberrant patterns of functional connectivity (FC), often manifesting as altered FC between the amygdala and PFC/ACC, are considered key features underlying the core symptom of worry.

This observed functional abnormality may reflect a structural impairment in the underlying communication infrastructure, i.e., the white matter (WM) tracts that provide anatomical connections between the PFC, ACC, and the limbic system (10). Diffusion tensor imaging (DTI), a noninvasive method for assessing WM microstructural integrity using metrics such as fractional anisotropy (FA), has been an indispensable tool for testing this structural hypothesis (6). A substantial body of evidence derived from DTI studies supports microstructural involvement, with numerous reports of reduced FA (indicative of decreased microstructural integrity) in key frontolimbic pathways (11, 12). Notably, microstructural deficits are often reported in the uncinate fasciculus (UF), a principal WM tract directly connecting the amygdala and frontal cortex (13, 14). Furthermore, WM alterations have been found in several other tracts implicated in anxiety-related networks, including the corpus callosum, cingulum, anterior thalamic radiation, inferior fronto-occipital fasciculus, inferior longitudinal fasciculus, and corona radiata (11, 15, 16). Importantly, the literature on structural network in GAD is not uniform; some studies even report focal increases in FA (6, 16, 17). This heterogeneity, encompassing both reduced and increased FA in different WM tracts, suggests that GAD may involve a complex and varying profile of WM alterations across the brain rather than a simple, uniform pattern of microstructural pathology. The lack of consistent findings on microstructural properties of WM tracts, even when accounting for the subtle effects noted by large-scale analyses such as meta- or mega-analysis, may also be due to significant methodological limitations in the prior research (6, 18).

Limitations from published DTI studies include the use of low-field (1.5T) or high-field (3T) MRI scanners, which provide inherently low signal-to-noise ratios (SNR), making it difficult to reliably detect subtle WM microstructural abnormalities. Further compounding the low SNR issue is the diverse DTI analytical approaches (e.g., region-based analysis or tract-based spatial statistics (19)) and high heterogeneity across study cohorts, stemming from small and mixed samples, varying comorbid conditions, differences in medication status, and broad age ranges. Together, these confounding factors make it difficult to isolate the specific pathology associated with GAD.

Here, we implemented a DTI with ultra-high field (7 Tesla, 7T), which offers superior resolution and sensitivity compared to lower-field systems (1.5T or 3T). The substantial increase in SNR and spatial resolution offered by 7T provides improved clarity. Instead of relying on predefined regions of interest (ROIs) for structural analysis of WM, we employed an advanced tractometry approach, i.e., correlational tractography, to delineate and investigate potential alterations in WM tract associated with GAD at the whole-brain level, allowing for comprehensive and fine-grained detection of microstructural changes that may have been overlooked in previous studies. Furthermore, we mitigated the confounding variables of chronic illness duration, aging effects, and medication exposure by selecting a sample of young adults with GAD. We hypothesized that patients with GAD would exhibit specific, dimensionally relevant alterations in diffusion-derived measures within the WM tracts critical to the circuitry of cortico-limbic emotion regulation, and these microstructural changes may correlate with the severity of dimensional measures of symptoms such as worry and anxiety. The goal of the present study is to generate robust, high-resolution evidence that helps refine and potentially reconcile existing competing neurobiological models, thereby advancing the field towards identifying reliable WM biomarkers for GAD.

## 2 Materials and Methods

### 2.1 Participants

A total of 88 participants were recruited in this study, including 27 patients with GAD and 61 healthy control participants (HCs), through the Pitt+Me portal by University of Pittsburgh Clinical and Translational Science Institute. GAD diagnoses were established by clinical psychiatrists according to the Diagnostic and Statistical Manual of Mental Disorders, Fifth Edition (DSM-V), and confirmed by experienced clinical psychologists through a medical record review process. Exclusion criteria for all participants across the groups included: (1) current or past major neurological or systemic medical illness; (2) current or past schizophrenia, bipolar disorder, post-traumatic stress disorder, mania, intellectual disabilities, obsessive-compulsive disorder, or social phobia; (3) history of alcohol abuse or dependence within the last 12 months; (4) self-reported history of substance abuse within the last 2 months; (5) contraindications to MRI (e.g., metallic implants, claustrophobia, or pregnancy); (6) the Wechsler Abbreviated Scale of Intelligence (Second Edition) < 70 or > 130; and (7) moderate or severe traumatic brain injury. Additionally, GAD participants with non-lifetime major depressive disorder (MDD) within the past month, treatment with high dose psychotropic medications or any antipsychotics, or present as a risk to themselves or others were also excluded. HCs were group-matched to the patient cohort by age, sex, and years of education. During the screening, research staff confirmed the absence of any current or past psychiatric disorders, and the exclusion criteria for all the participants across the groups were applied, with the additional requirement of no family history of psychiatric illness. The study protocol was approved by the Institutional Review Board (IRB) of the University of Pittsburgh and conducted in accordance with the Declaration of Helsinki. Written informed consent was obtained from all participants prior to enrollment. Parents or legal guardians provided assistance to adolescent participants in understanding the study procedures and consent/assent documents, consistent with IRB guidelines for obtaining informed permission and assent from minors. During clinical assessment, all participants completed the Hamilton Anxiety Rating Scale (HAM-A) and Penn State Worry Questionnaire (PSWQ) to quantify anxiety and worry severity.

### 2.2 MRI Data Acquisition and Processing

Diffusion-weighted images were acquired on a 7T human MRI scanner (Siemens Magnetom, Erlangen, Germany) with a custom-built 60 Tx/32Rx Tic-Tac-Toe (TTT) head radiofrequency (RF) coil system (20, 21). Specifically, we employed a single-shell DTI sequence with following imaging parameters; repetition time (TR) = 6500 ms, echo time (TE) = 81.6 ms, number of b_0_ = 4, diffusion sampling directions = 46, *b*-value = 1010 s/mm², in-plane resolution = 1.5 mm, and slice thickness = 1.5 mm. Preprocessing of the diffusion-weighted imaging data began with a bias correction step to mitigate signal inhomogeneity (22). Next, the susceptibility artifact was estimated using reversed phase-encoding b_0_ images processed with TOPUP in the TinyFSL package (https://fsl.fmrib.ox.ac.uk/fsl/). Subsequently, eddy current and head movement corrections were performed using FSL’s EDDY, implemented through the integrated preprocessing interface in DSI Studio (23). The accuracy of the b-table orientation was then validated by comparing fiber orientations against a population-averaged template (24). Finally, the corrected DWI data were reconstructed in the standard Montreal Neurological Institute (MNI) space using *q*-space diffeomorphic reconstruction (QSDR) that nonlinearly transforms the images to the ICBM152 adult template (25). This process yielded the spin distribution function using a diffusion sampling length ratio of 1.25 and an isotropic output resolution of 1.5 mm (26). The diffusion scalar indices, including FA and mean diffusivity (MD), were estimated for use in the connectometry analysis.

### 2.3 Correlational Tractography

Diffusion MRI connectometry analysis was performed to identify WM pathways using correlational tractography in which local diffusion indices (i.e., FA and MD) were correlated with group differences (27). A nonparametric Spearman partial correlation was used to assess voxel-wise associations between changes in diffusion indices along fiber pathways and group while controlling for age, sex, years of education, handedness, and size of lateral ventricle using multiple regression. Controlling for lateral ventricle volume was intended to reduce the influence of partial volume effects at the ventricle-WM interface. The analysis included all 88 participants. A *t*-score threshold of 2.5 was applied, and correlational tractography was conducted using a deterministic fiber tracking algorithm (28). The cerebellum was excluded due to signal variation across subjects, and whole-brain seeding was implemented at MNI coordinates. The resulting tracts were refined using topology-informed pruning with 16 iterations (29). To correct for multiple comparisons, a false discovery rate (FDR) threshold of 0.05 was applied, and statistical significance was determined by estimating the null distribution of track length through 4,000 randomized permutations of the group labels. The coordinates of the identified WM tract bundles were saved and subsequently used to extract local diffusion indices for statistical analysis.

### 2.4 Neuropsychological Assessments

To assess anxiety severity and worry-related symptoms, all participants completed the HAM-A and PSWQ. The HAM-A (30) is a clinician-administered instrument designed to evaluate the severity of both psychic and somatic symptoms of anxiety. It comprises 14 items, each rated on a 5-point scale from 0 (absent) to 4 (very severe), yielding total scores ranging from 0 to 56, with higher scores indicating greater anxiety severity. Scores of 0–17 indicate no-to-mild anxiety, 18–24 moderate anxiety, and 25–30 severe anxiety. The PSWQ (31) is a self-report measure designed to assess the tendency toward excessive and uncontrollable worry, the hallmark feature of GAD. It consists of 16 items, each rated from 1 (not at all typical of me) to 5 (extremely typical of me), producing a total score ranging from 16 to 80, with higher scores reflecting greater psychological worry. We used the total scores from both the HAM-A and PSWQ to assess the overall anxiety severity and trait worry in our cohort.

### 2.5 Statistical Analysis

We first performed a confirmatory group comparison using analyses of covariance (ANCOVAs) adjusted for age, sex, years of education, handedness, and lateral ventricle volume. Multiple comparisons were corrected using the Benjamini-Hochberg method, accounting for the number of identified tract bundles. Because each tract bundle comprised streamlines extending across multiple regions, we further examined spatial patterns of WM alterations by dividing the identified bundles into three main segments along their principal orientations. Separate ANCOVAs were then conducted for each segment with the same covariates, and multiple comparison correction was applied to control for alpha inflation.

To examine clinical associations of altered WM bundles, we performed linear regression analyses with robust estimation to test whether diffusion indices of the identified tracts could explain variance in symptom severity as measured by total scores on the HAM-A and PSWQ while adjusting for the same covariates used in the ANCOVAs. *Post hoc* analyses were also conducted on individual tract segments to evaluate the spatial pattern of effect sizes using partial correlation, with Benjamini-Hochberg correction applied to control for multiple comparisons.

Finally, to assess the potential of the identified WM tracts as diagnostic-aided markers, we conducted a classification analysis to differentiate patients with GAD from HCs using both symptom severity scores and tract-based diffusion indices. Total HAM-A and PSWQ scores were used as indicators of symptom severity. Four support vector machine (SVM) classifiers were trained (32); one for each neuropsychological score, one for the WM features, and one combined model integrating all features. Before model training, all input features were adjusted for the effects of covariates. The SVMs were implemented with a cubic kernel and evaluated using receiver operating characteristic (ROC) analysis and performance metrics including accuracy, sensitivity, and specificity, based on five-fold cross-validation.

## 3 Results

### 3.1 Characteristics of study cohorts

The basic demographic and neuropsychological characteristics of the study cohorts are shown in Table 1. There were no statistical differences in terms of age (*P* = 0.800), sex (*P* = 0.731), education (*P* = 0.584), and handedness (*P* = 0.670). The patients with GAD showed significantly higher scores of HAM-A and PSWQ than the HCs (*P’s* < 0.001).

**Table 1.** Characteristics of the participants in the study.

| Characteristics | GAD | HC | Comparisons |
| --- | --- | --- | --- |
| N | 27 | 61 | - |
| Age, year | 21.9 (2.5) | 21.7 (2.4) | $P = 0.800$ |
| Sex, female, N (%) | 20 (74.1%) | 43 (70.5%) | $P = 0.731$ |
| Education, year | 15.5 (2.0) | 15.2 (2.0) | $P = 0.584$ |
| Handedness, right, N (%) | 24 (88.9%) | 50 (82.0%) | $P = 0.670$ |
| PSWQ total score | 62.7 (12.5) | 35.9 (11.6) | $P < 0.001$ |
| HAM-A total score | 24.3 (9.2) | 4.7 (4.8) | $P < 0.001$ |

Note: Continuous and categorical variables were tested using two-sample $t$ test and chi-square test, respectively. Abbreviation: GAD, Generalized Anxiety Disorder; HC, Healthy Control; PSWQ, Penn State Worry Questionnaire; HAM-A, Hamilton Anxiety Rating Scale.

### 3.2 Distinct alterations of WM tract bundles in GAD

Whole brain FDR-corrected correlational tractography identified two fiber tract bundles showing significantly altered FA associated with GAD. The confirmatory group comparison indicated that one tract bundle connecting the right limbic region to the right PFC exhibited significantly decreased FA in patients with GAD compared to the HCs (GAD: 0.303 ± 0.096, HCs: 0.352 ± 0.098, *F*_(1,81)_ = 4.40, *P_FDR_* = 0.039) (Figure 1a). In contrast, another tract bundle within the right cingulum demonstrated significantly increased FA in patients with GAD relative to HCs (GAD: 0.491 ± 0.060, HCs: 0.401 ± 0.084, *F*_(1,81)_ = 22.74, *P_FDR_* < 0.001) (Figure 1b). These two identified tract bundles were not significantly correlated with each other according to the partial correlation analysis. (*rho* = -0.119, *P* = 0.285). On the other hand, the correlational tractography analysis did not identify any WM tract bundles showing significant associations with GAD when based on MD maps.

**Figure 1.**
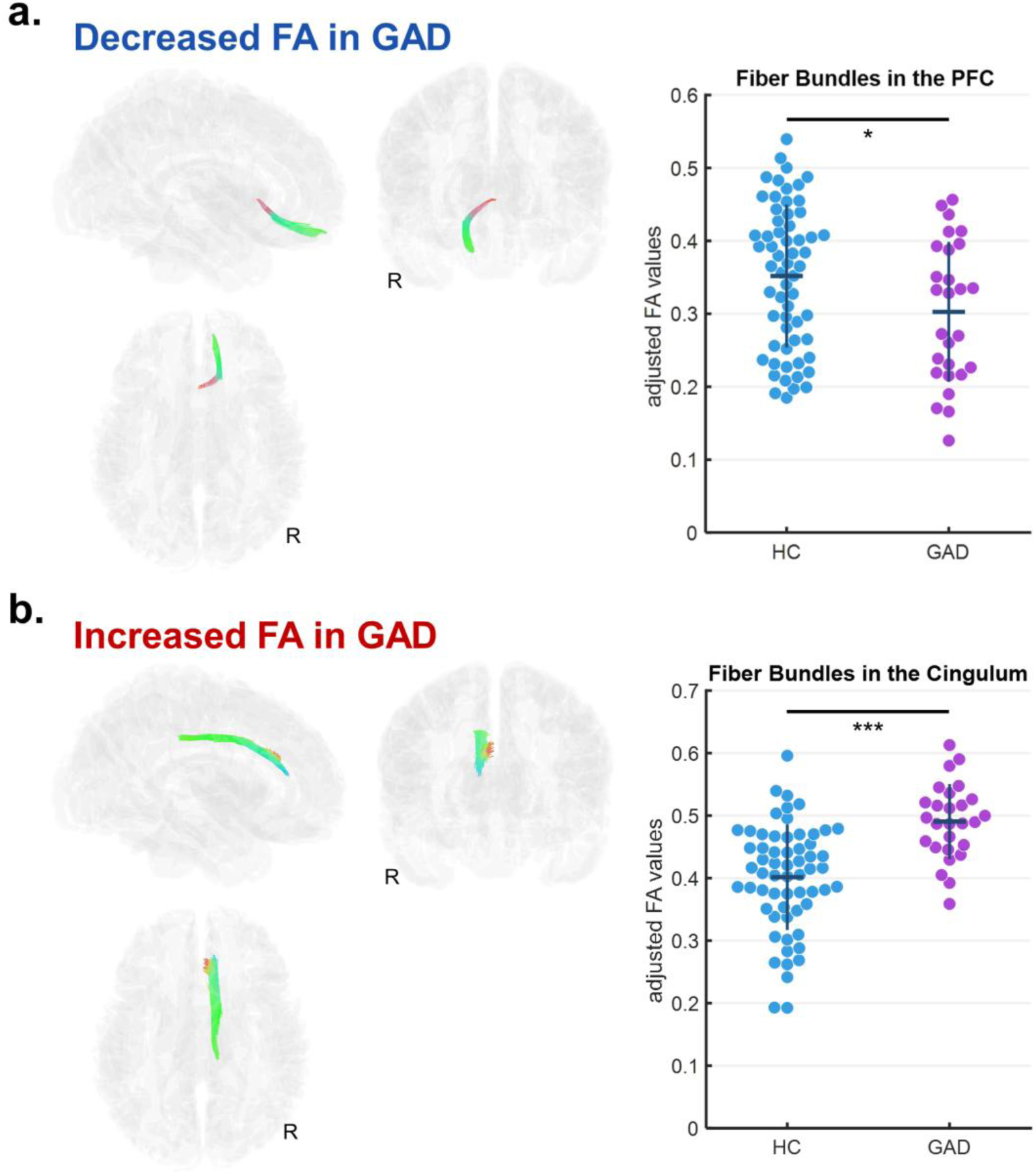
Correlational tractography of WM pathways associated with GAD based on FA. Two distinct patterns of altered connectivity were observed: (a) fiber bundles connecting the right limbic region to the right PFC showed significantly decreased FA in the patients with GAD, whereas (b) fiber bundles within the right cingulum demonstrated significantly increased FA in the patients with GAD. Analyses controlled for age, sex, education, handedness, and lateral ventricle volume. The tract bundles are color-coded to indicate orientation; green, red, and blue represent the anterior-posterior, left-right, and superior-inferior directions, respectively. Abbreviation: FA, fractional anisotropy; GAD, generalized anxiety disorder; PFC, prefrontal cortex; WM, white matter.

For the tract bundles showing significantly altered FA in GAD, we conducted segment-wise group comparisons along their principal orientations. As both identified bundles were association fibers extending along the anterior-posterior axis, each fiber bundle was divided into anterior, middle, and posterior segments. In the tract connecting to the right PFC, the posterior and middle segments showed no significant group differences (posterior region, GAD: 0.302 ± 0.105, HCs: 0.333 ± 0.103, *F*_(1,81)_ = 1.56, *P_FDR_* = 0.215; middle region, GAD: 0.336 ± 0.116, HCs: 0.385 ± 0.114, *F*_(1,81)_ = 3.23, *P_FDR_* = 0.114) whereas the anterior segment demonstrated a significant reduction in FA in patients with GAD compared to HCs (anterior region, GAD: 0.293 ± 0.124, HCs: 0.362 ± 0.115, *F*_(1,81)_ = 6.05, *P_FDR_* = 0.048) (Figure 2a). The effect size increased progressively from the posterior to the anterior segments, suggesting that FA reduction was more pronounced toward the prefrontal region in GAD. In contrast, the tract within the right cingulum exhibited significant group differences across all segments (posterior region, GAD: 0.563 ± 0.062, HCs: 0.490 ± 0.093, *F*_(1,81)_ = 12.79, *P_FDR_* < 0.001; middle region, GAD: 0.527 ± 0.081, HCs: 0.422 ± 0.107, *F*_(1,81)_ = 19.40, *P_FDR_* < 0.001; anterior region, GAD: 0.413 ± 0.063, HCs: 0.312 ± 0.092, *F*_(1,81)_ = 24.82, *P_FDR_* < 0.001), with patients with GAD showing higher FA values, particularly in the middle and anterior portion (Figure 2b). Similarly, effect sizes increased toward the anterior (frontal) region, indicating a gradient of more prominent FA elevation in the frontal segments of the right cingulum.

**Figure 2.**
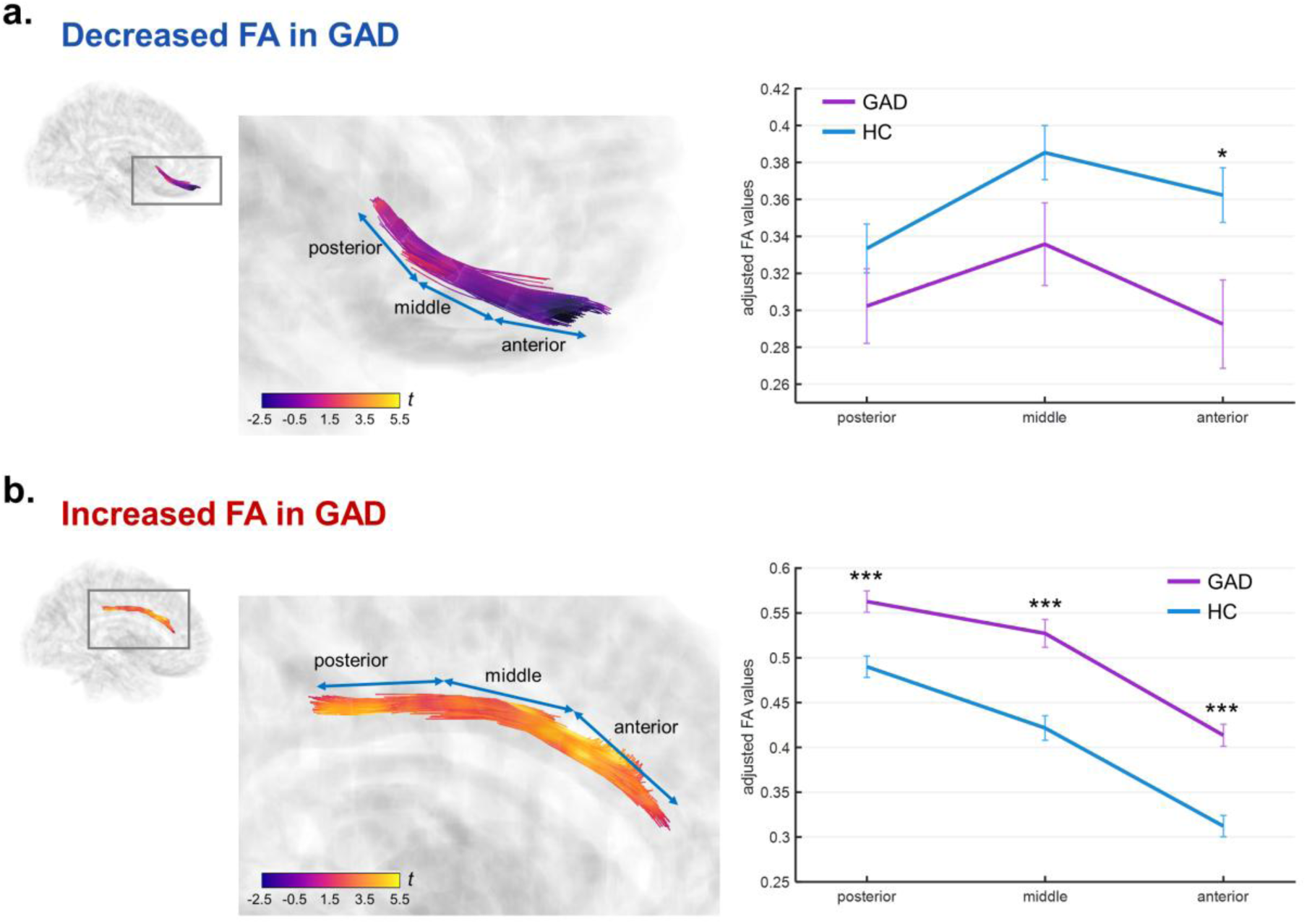
Segment-wise group comparisons of FA along the identified WM tracts. Each tract was divided into anterior, middle, and posterior segments along the anterior–posterior axis. (a) The tract connecting the right limbic region to the right PFC showed a significant FA reduction in patients with GAD compared to healthy controls, primarily in the anterior segment. (b) The right cingulum tract exhibited significantly higher FA in GAD across all segments, with the effect most pronounced in the anterior (frontal) region. Analyses controlled for age, sex, education, handedness, and lateral ventricle volume. The tract bundles are color-coded to represent the corresponding statistical *t*-values. Abbreviation: FA, fractional anisotropy; GAD, generalized anxiety disorder; WM, white matter.

### 3.3 Clinical associations of identified WM tract bundles with symptom severity

We investigated whether the identified tract bundles were associated with clinical symptom severity measured by the HAM-A for general/overall anxiety and PSWQ for trait worry. Regression analyses showed that FA values in the prefrontal tract bundle were not significantly correlated with either PSWQ or HAM-A scores (PSWQ, β = -32.5, standard error [SE] = 20.5, 95% confidence interval [C.I.] = [-73.3, 8.4], *P_FDR_* = 0.117; HAM-A, β = -21.6, SE = 12.2, 95% C.I. = [-45.8, 2.6], *P_FDR_* = 0.107) (Figures 3a and 3c). In contrast, FA values in the right cingulum tract exhibited significant positive associations with both measures (PSWQ, β = 98.1, SE = 21.7, 95% C.I. = [54.9, 141.4], *P_FDR_* < 0.001; HAM-A, β = 49.1, SE = 12.6, 95% C.I. = [23.9, 74.2], *P_FDR_* < 0.001) (Figures 3b and 3d), indicating that higher FA values were linked to greater trait worry- and general anxiety-related symptom severity.

**Figure 3.**
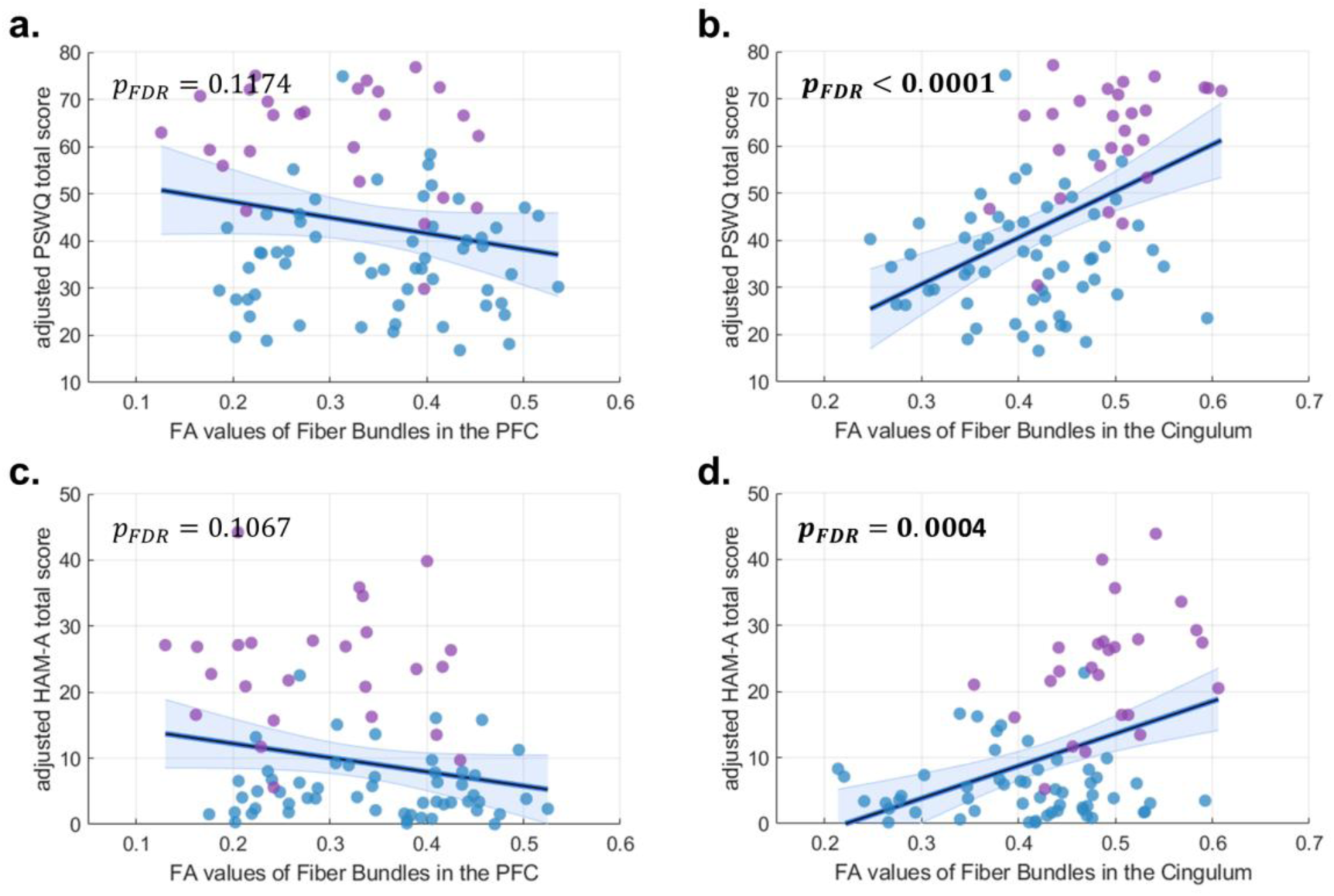
Associations between FA in the identified WM tracts and clinical symptom severity. Panels (a) and (c) show that FA values of tract bundle in the right PFC were not significantly correlated with PSWQ or HAM-A total scores. Panels (b) and (d) illustrate that FA values in the right cingulum tract showed significant positive associations with both PSWQ and HAM-A scores, indicating that higher FA values were linked to greater trait worry- and general anxiety-related symptom severity. Analyses controlled for age, sex, education, handedness, and lateral ventricle volume. Purple and blue colors indicate patients with GAD and HCs, respectively. Abbreviation: FA, fractional anisotropy; GAD, generalized anxiety disorder; HAM-A, Hamilton Anxiety Rating Scale; HC, healthy control; PFC, prefrontal cortex; PSWQ, Penn State Worry Questionnaire.

Similarly, we examined the segment-wise associations of the right cingulum tract with clinical symptom severity. For PSWQ, the middle segment showed the strongest partial correlation (*rho* = 0.454, 95% C.I. = [0.435, 0.473], *P_FDR_* < 0.001), followed by the anterior (*rho* = 0.416, 95% C.I. = [0.395, 0.436], *P_FDR_* < 0.001) and posterior (*rho* = 0.346, 95% C.I. = [0.324, 0.367], *P_FDR_* = 0.005) segments (Figure 4a). For HAM-A, the anterior segment exhibited the greatest positive correlation (*rho* = 0.439, 95% C.I. = [0.419, 0.458], *P_FDR_* < 0.001), followed by the middle (*rho* = 0.424, 95% C.I. = [0.404, 0.443], *P_FDR_* < 0.001) and posterior (*rho* = 0.298, 95% C.I. = [0.276, 0.320], *P_FDR_* = 0.023) segments (Figure 4b). These findings align with our earlier observation that the anterior portion of the right cingulum (close to the frontal region) shows a stronger effect in GAD compared to the more posterior segments.

**Figure 4.**
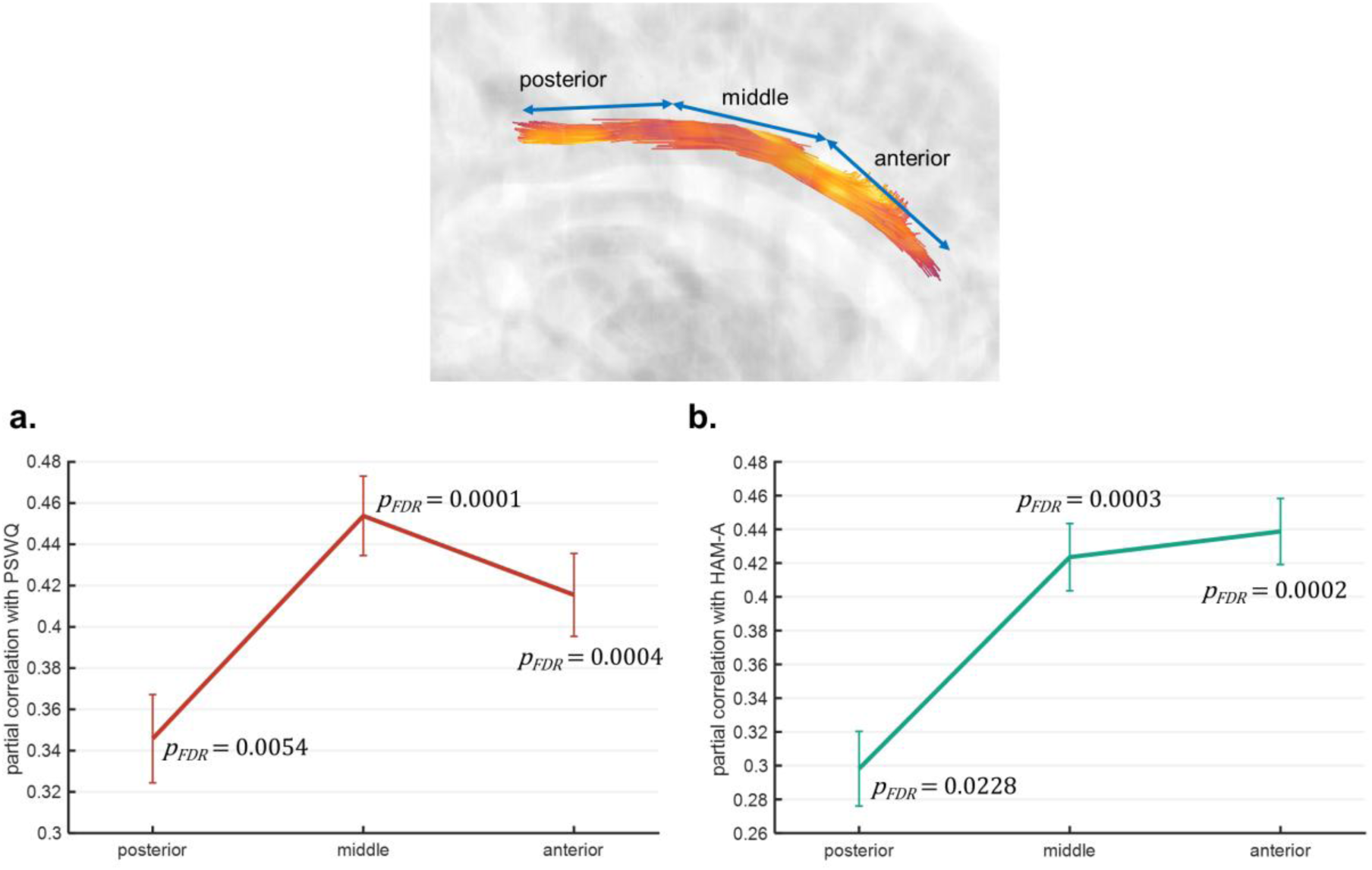
Segment-wise associations between FA in the right cingulum and clinical symptom severity. Panels (a) and (b) indicate the partial correlation coefficients of PSWQ and HAM-A with FA values across segments, respectively. Analyses controlled for age, sex, education, handedness, and lateral ventricle volume. Error bars represent 95% confidence intervals of the estimates. Abbreviation: FA, fractional anisotropy; HAM-A, Hamilton Anxiety Rating Scale; PSWQ, Penn State Worry Questionnaire.

### 3.4 Improvement of classification between GAD and HC using identified WM tract bundles

To evaluate the clinical value of the altered WM tract bundles as a diagnostic-aided marker, we conducted a classification task to differentiate GAD and HC groups by using different combinations of neuropsychological scores (i.e., PSWQ & HAM-A) as well as WM features and assessed the performance by using receiver operating characteristic (ROC) analysis. By using the SVM modeling with a cubic kernel through 5-fold cross-validation, the baseline model using PSWQ achieved an accuracy of 85.2% (95% C.I. = [85.0%, 85.5%]) and area under the ROC curve (AUC) = 0.876 (95% C.I. = [0.873, 0.879]) (Figure 5). The sensitivity and specificity were 0.630 (95% C.I. = [0.624, 0.635]) and 0.951 (95% C.I. = [0.949, 0.953]), respectively. By using HAM-A, the performance was 75.0% (95% C.I. = [74.7%, 75.3%]) in accuracy and 0.735 (95% C.I. = [0.731, 0.739]) in AUC. The sensitivity and specificity were 0.704 (95% C.I. = [0.698, 0.709]) and 0.770 (95% C.I. = [0.767, 0.774]), respectively. When combining both neuropsychological scores, the performance can be enhanced; accuracy = 87.5% (95% C.I. = [87.3%, 87.7%]), AUC = 0.910 (95% C.I. = [0.908, 0.913]), sensitivity = 0.815 (95% C.I. = [0.810, 0.819]), and specificity = 0.902 (95% C.I. = [0.899, 0.904]). The classification performance using only WM features (i.e., FA values from the right prefrontal and cingulum tracts) was relatively lower than those achieved with neuropsychological measures; accuracy = 70.5% (95% C.I. = [70.1%, 70.8%]), AUC = 0.724 (95% C.I. = [0.720, 0.727]), sensitivity = 0.407 (95% C.I. = [0.401, 0.413]), and specificity = 0.836 (95% C.I. = [0.833, 0.839]). However, the overall classification performance can be further improved by including WM features in the all-in-one model (Figure 5). The final model achieved an accuracy of 93.2% (95% C.I. = [93.0%, 93.3%]) and AUC = 0.967 (95% C.I. = [0.966, 0.968]), with high sensitivity = 0.852 (95% C.I. = [0.848, 0.856]) and high specificity = 0.967 (95% C.I. = [0.966, 0.968]). This result suggested that the identified WM tract features could provide additional information to enhance the differentiation between GAD and HCs. The coordinates of the identified tracts are available upon request from the corresponding author.

**Figure 5.**
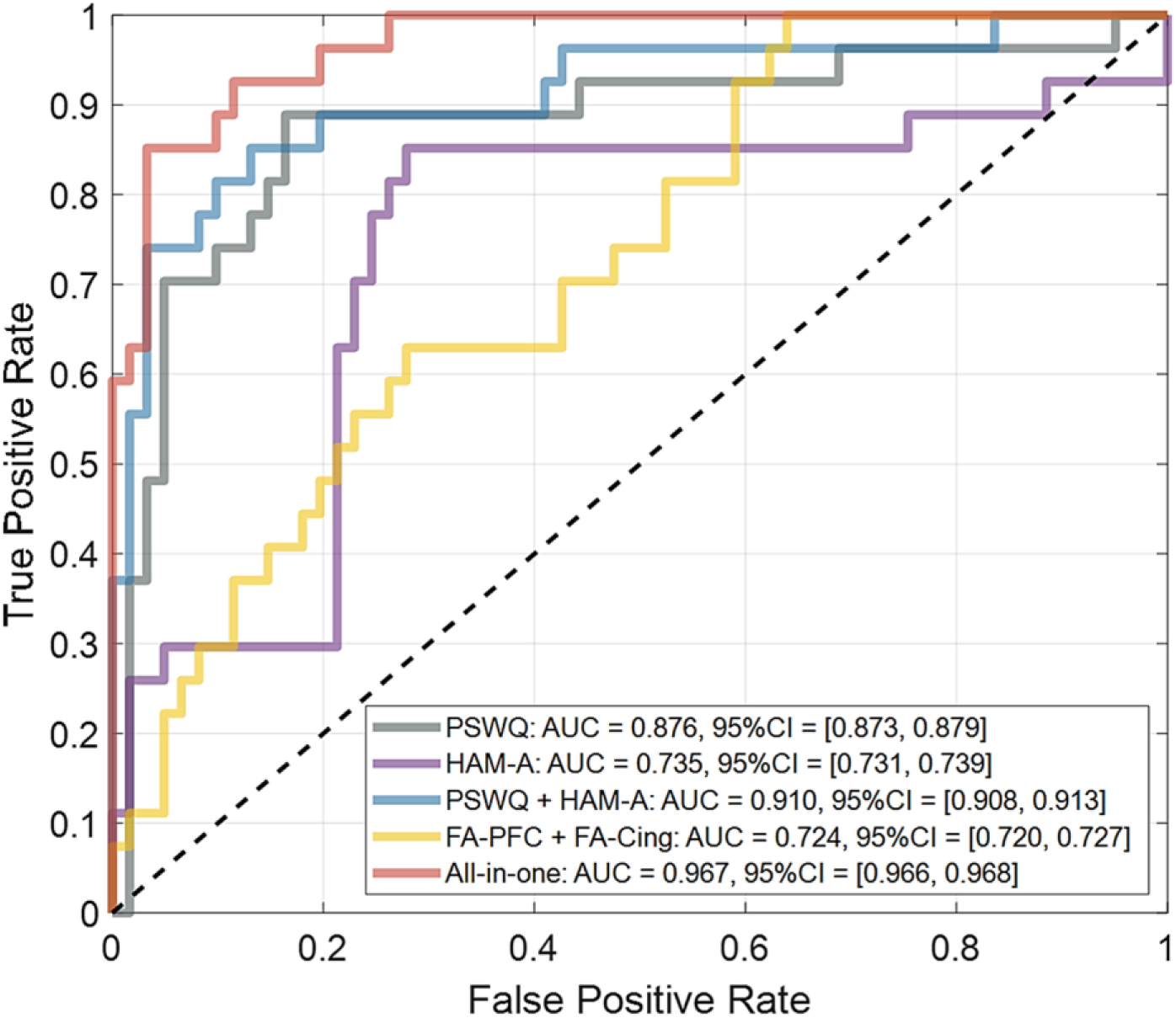
ROC analysis for GAD classification based on neuropsychological scores and identified WM tract features. The ROC curves were yielded by using different combinations of neuropsychological scores and/or WM features. The performance was evaluated by using 5-fold cross-validation. Abbreviation: AUC, area under curve; CI, confidence interval; Cing, cingulum; FA, fractional anisotropy; HAM-A, Hamilton Anxiety Rating Scale; PFC, prefrontal cortex; PSWQ, Penn State Worry Questionnaire; ROC, receiver operating characteristic.

## 4 Discussion

The present study employed ultra-high field 7T diffusion tensor imaging and correlational tractography to delineate fine-grained WM alterations in individuals with GAD compared with HCs. Two distinct, spatially dissociable tract abnormalities were identified: (1) decreased FA in fibers connecting the right limbic region and the right PFC, and (2) increased FA in the right cingulum. Notably, the most prominent alterations were observed in the anterior portions of both tracts, implicating prefrontal-limbic circuitry as a central structural substrate of anxiety pathophysiology. Furthermore, FA in the right cingulum was positively correlated with both anxiety and worry severity, and inclusion of these WM features improved diagnostic classification performance. These findings provide convergent structural evidence supporting distinct forms of altered fronto-limbic connectivity in emotion dysregulation models in GAD and demonstrate the higher sensitivity of the 7T tractometry for identifying differential microstructural changes that may not be detectable at lower field strengths.

Our finding of reduced FA in the prefrontal tract aligns with extensive evidence implicating compromised structural integrity along fronto-limbic pathways in GAD (11, 12). The uncinate fasciculus, a tract connecting the amygdala and ventromedial PFC, has consistently shown decreased FA in both adult and adolescent GAD cohorts (11, 13). This tract mediates top-down regulation of limbic responses and facilitates cognitive control of affective states (14, 33). Our finding of reduced FA toward the anterior segment suggests that diminished integrity in prefrontal terminations may specifically weaken regulatory input to subcortical emotion-processing regions (9, 14, 34). Functionally, such disruption may underlie the hypoactivation of the ventromedial and dorsolateral PFC that is frequently reported during emotion regulation and threat anticipation in GAD (3, 4). The anterior gradient of FA reduction further supports the notion that cortical dysregulation in GAD is driven primarily by compromised prefrontal connectivity rather than by primary limbic abnormalities (2). The observed reduction in FA may reflect reduced axonal coherence or altered fiber organization, leading to less efficient communication within the emotion regulation network (35, 36). This interpretation is consistent with functional MRI studies showing attenuated inhibitory regulation from the PFC to the amygdala, coupled with hyperactivation of salience-processing regions such as the ACC and insula (3, 4, 6). Although decreased FA in the prefrontal region did not show a significant association with worry or anxiety, the observed trend toward a negative correlation suggests a potential link between reduced WM integrity and greater symptom severity. These findings support a model in which weakened prefrontal WM connectivity compromises top-down control, promoting exaggerated limbic reactivity and sustained worry, the core clinical features of GAD.

In contrast, the right cingulum exhibited increased FA in GAD compared with HCs, most prominently in its anterior and middle segments, and these increases were positively correlated with both worry and anxiety scores. Elevated FA has been reported in several anxiety-related conditions (35, 37), and may reflect maladaptive compensatory WM reorganization rather than genuinely enhanced connectivity (6). The cingulum is a major associative tract interlinking the medial PFC, parietal cortex, and hippocampal regions, which are core components of the default mode and limbic networks that support self-referential thought and emotion regulation (3, 6). Heightened FA in this region may indicate enhanced or overly rigid synchronization within default mode circuits, consistent with the excessive self-focused and future-oriented rumination that characterizes GAD (6, 10). Functionally, anterior cingulate regions show aberrant neural activation during threat anticipation and the regulation of emotional conflict in GAD (3, 5, 6). These divergent findings are consistent with our structural MRI results: greater cingulum FA may reflect increased cingulate engagement in worry-related processing. This interpretation is further supported by our segment-wise correlation analyses, which show that FA in anterior cingulum, proximal to prefrontal targets, was strongly associated with worry- and anxiety-related symptom severity.

In the present study, we demonstrated the capability of 7T diffusion-weighted imaging combined with correlational tractography to resolve fine-scale WM alterations that are challenging to detect using lower-field (e.g., 1.5T or 3T) scanners. The superior SNR and spatial resolution of 7T enabled the identification of opposing microstructural changes in neighboring tracts, thereby revealing the differential nature of WM pathology in GAD. In contrast to traditional predefined region- or tract-based approaches (38), our whole-brain tractometry method captured spatially continuous and segment-specific variations in diffusion metrics, revealing their continuous, along-tract gradients. This analytic framework may help reconcile prior inconsistencies in the DTI literature, where findings often varied with how ROIs were selected. Integration of WM features into the machine-learning classifier further underscores their potential for clinical translation (39). While neuropsychological measures alone yielded relatively high diagnostic accuracy, incorporating diffusion MRI-derived indices enhanced classification performance beyond behavioral data. This suggests that structural connectivity metrics may serve as complementary markers for individualized assessment of anxiety severity and treatment response, especially as imaging-based tools become increasingly feasible and practical in clinical research.

Several limitations merit consideration. First, the cross-sectional design precludes inferences about causality and characterization of the temporal evolution of WM changes. Longitudinal studies are needed to clarify whether the observed microstructural differences reflect premorbid vulnerability, compensatory adaptation, or chronic sequelae of anxiety. Second, although the sample was restricted to an age range from 16 to 26 years and was imbalanced across study cohorts, as new onset of GAD increases substantially from this age range, we may need to expand the age range to 10-30 years old and increase sample size to capture a more complete trajectory of WM changes during the sensitive development period of anxiety-related disorders. Third, reliance on FA, a sensitive yet nonspecific measure of microstructural integrity, should be complemented by additional diffusion metrics (e.g., neurite density, free-water imaging) and multimodal integration of functional and metabolic data would sharpen mechanistic interpretations (4).

In brief, this 7T DTI study reveals a differential pattern of WM alterations in GAD, characterized by reduced FA in right prefrontal tracts and increased FA in the right cingulum, both exhibiting an anterior predominance. These opposing microstructural changes likely reflect distinct facets of the disorder but collectively provide a structural framework for understanding the persistent worry and emotional dysregulation that are central to GAD. By integrating ultra-high field tractometry with symptom-dimensional analysis, our findings help refine the current neurobiological model of GAD and underscore the potential of WM metrics as imaging markers for early detection and treatment stratification in anxiety disorders.

## 5 Acknowledgements

This work was supported by the National Institute of Mental Health (NIMH) with grant; R21-MH116475.

## 8 Author’s Contribution

(1) Research Project: A. Conception, B. Organization, C. Execution; (2) Statistical Analysis: A. Design, B. Execution, C. Review and Critique; (3) Manuscript Preparation: A. Writing of the First Draft, B. Review and Critique.

C.L. Chen: 1A, 1B. 1C, 2A, 2B, 3A; M. Wu: 1A, 2A, 2C, 3B; Y. Chi: 1B, 1C; M. Rose: 1C, 3B; J.C. Weber: 1C, 3B; S.T. Lamb: 1C, 3B; H. Jin: 2C, 3B; T.S. Ibrahim: 1C, 3B; C.D. Ladouceur: 1A, 3B; Y.F. Chang: 1A, 2C, 3B; K.Y. Tseng: 1A, 2C, 3B; H.J. Aizenstein: 1A, 2A, 2C, 3B; S. Yang: 1A, 2A, 2C, 3A, 3B.

## 9 Consent Statement

All procedures performed in this study involving human participants from the FANSDA (Functional and Neurochemical Substrates of Amygdala-Frontal Circuitry across Development and Anxiety) project were in accordance with the ethical standards of the Institutional Review Boards of University of Pittsburgh Medical Center (no.STUDY21020052) and with the 1964 Helsinki declaration and its later amendments or comparable ethical standards. Informed consent in the study was obtained from all individual participants who were recruited in this project.

